# Multiethnic meta-analysis identifies new loci for pulmonary function

**DOI:** 10.1101/196048

**Authors:** Annah B. Wyss, Tamar Sofer, Mi Kyeong Lee, Natalie Terzikhan, Jennifer N. Nguyen, Lies Lahousse, Jeanne C. Latourelle, Albert Vernon Smith, Traci M. Bartz, Mary F. Feitosa, Wei Gao, Tarunveer S. Ahluwalia, Wenbo Tang, Christopher Oldmeadow, Qing Duan, Kim de Jong, Mary K. Wojczynski, Xin-Qun Wang, Raymond Noordam, Fernando Pires Hartwig, Victoria E. Jackson, Tianyuan Wang, Ma’en Obeidat, Brian D. Hobbs, Tianxiao Huan, Gleb Kichaev, Jianping Jin, Mariaelisa Graff, Tamara B. Harris, Ravi Kalhan, Susan R. Heckbert, Lavinia Paternoster, Kristin M. Burkart, Yongmei Liu, Elizabeth G. Holliday, James G. Wilson, Judith M. Vonk, Jason Sanders, R. Graham Barr, Renée de Mutsert, Ana Maria Baptista Menezes, Hieab H. H. Adams, Maarten van den Berge, Roby Joehanes, Lenore J. Launer, Alanna C. Morrison, Colleen M. Sitlani, Juan C. Celedón, Stephen B. Kritchevsky, Rodney J. Scott, Kaare Christensen, Jerome I. Rotter, Tobias N. Bonten, Fernando César Wehrmeister, Yohan Bossé, Nora Franceschini, Jennifer A. Brody, Robert C. Kaplan, Kurt Lohman, Mark McEvoy, Michael A. Province, Frits R. Rosendaal, Kent D. Taylor, David C. Nickle, Kari E. North, Myriam Fornage, Bruce M. Psaty, Richard H. Myers, George O’Connor, Torben Hansen, Cathy C. Laurie, Pat Cassano, Joohon Sung, Woo Jin Kim, John R. Attia, Leslie Lange, H. Marike Boezen, Bharat Thyagarajan, Stephen S. Rich, Dennis O. Mook-Kanamori, Bernardo Lessa Horta, André G Uitterlinden, Don D. Sin, Hae Kyung Im, Michael H. Cho, Guy G. Brusselle, Sina A. Gharib, Josée Dupuis, Ani Manichaikul, Stephanie J. London

## Abstract

Nearly 100 loci have been identified for pulmonary function, almost exclusively in studies of European ancestry populations. We extend previous research by meta-analyzing genome-wide association studies of 1000 Genomes imputed variants in relation to pulmonary function in a multiethnic population of 90,715 individuals of European (N=60,552), African (N=8,429), Asian (N=9,959), and Hispanic/Latino (N=11,775) ethnicities. We identified over 50 novel loci at genome-wide significance in ancestry-specific and/or multiethnic meta-analyses. Recent fine mapping methods incorporating functional annotation, gene expression, and/or differences in linkage disequilibrium between ethnicities identified potential causal variants and genes at known and newly identified loci. Sixteen of the novel genes encode proteins with predicted or established drug targets, including *KCNK2* and *CDK12.*

## Introduction

Pulmonary function traits (PFTs), including forced expiratory volume in the first second (FEV_1_) and forced vital capacity (FVC), and their ratio FEV_1_/FVC, are important clinical measures for assessing the health of the lungs, diagnosing chronic obstructive pulmonary disease (COPD), and monitoring the progression and severity of various other lung conditions. Further, even when within the normal range, these parameters are related to mortality, independently of standard risk factors^1-3^.

In addition to lifestyle and environmental factors, such as smoking and air pollution, there is a demonstrated genetic component to pulmonary function^4-6^. Previous genome-wide association studies (GWAS) have collectively identified nearly 100 loci associated with PFTs. These analyses have been primarily conducted using HapMap imputed data among European ancestry populations^7-12^. Recently, the UK BiLEVE Study (N=48,943) and SpiroMeta Consortium (N=38,199) have also examined associations between 1,000 Genomes imputed variants and PFTs, but only among Europeans^13-15^.

The present <u>C</u>ohorts for <u>H</u>eart and <u>A</u>ging <u>R</u>esearch in <u>G</u>enomic <u>E</u>pidemiology (CHARGE) meta-analysis builds upon previous studies by examining PFTs in relation to the more comprehensive 1000 Genomes panel in a larger study population (90,715 individuals from 22 studies, Table 1) comprised of multiple ancestral populations: European (60,552 individuals from 18 studies), African (8,429 individuals from 7 studies), Asian (9,959 individuals from 2 studies), and Hispanic/Latino (11,775 individuals from 6 ethnic background groups in 1 study). Along with look-up of our top findings in existing analyses of lung function traits and COPD, we additionally investigated correlation with gene expression in datasets from blood and lung tissue and assessed the potential biological and clinical relevance of our findings using recently developed methods integrating linkage disequilibrium (LD), functional annotation, gene expression and the multiethnic nature of our data. Finally, we identified known or potential drug targets for newly identified lung function associated loci.

**Table 1.**
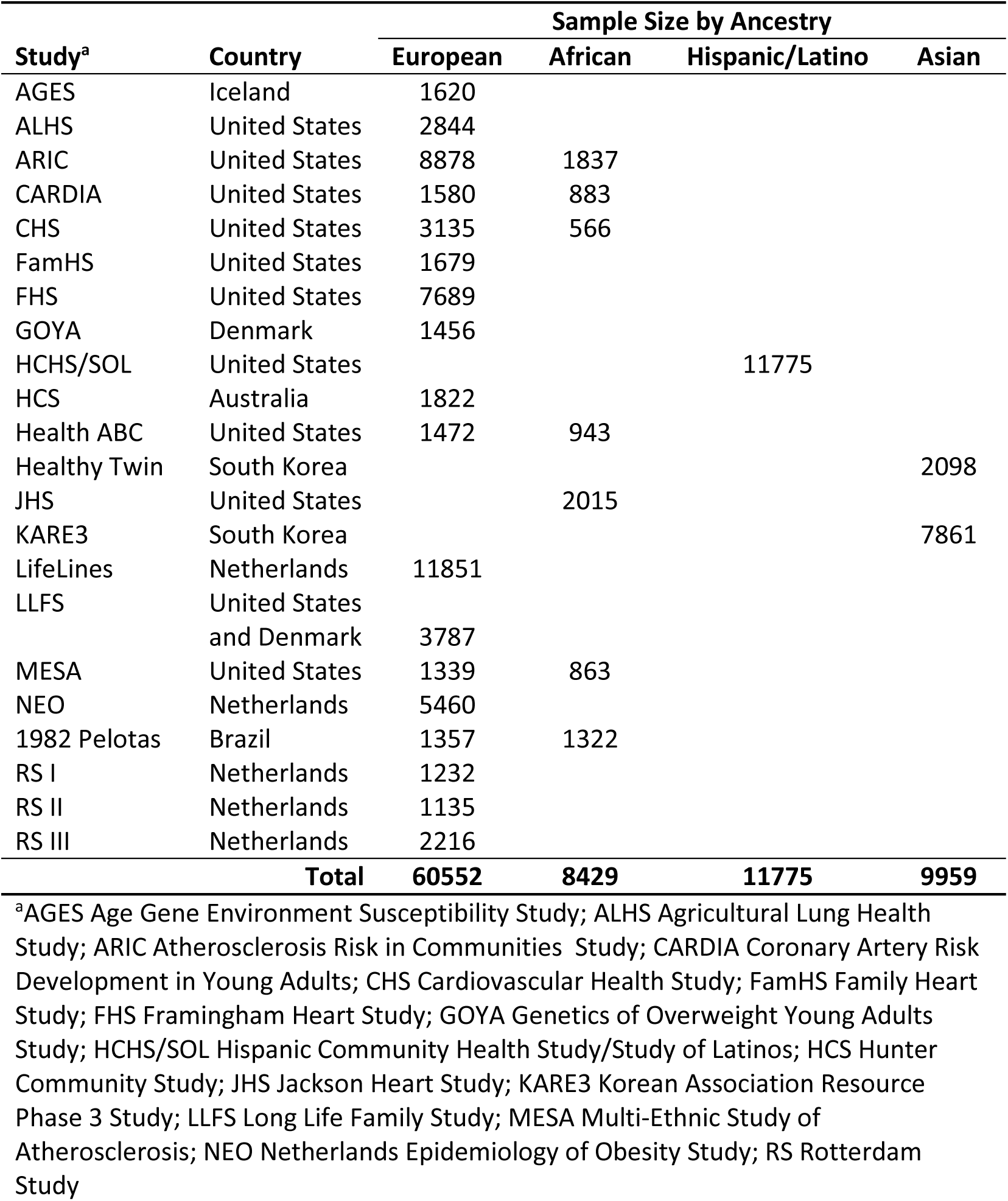
Sample size and location of studies included in the CHARGE consortium 1000 Genomes and pulmonary function meta-analysis

## Results

### Ancestry-Specific Meta-Analyses

Each study used linear regression to model the additive effect of variants on PFTs, adjusting for age, sex, height, cigarette smoking, center (if multicenter study), and ancestral principal components, including a random familial effect to account for family relatedness when appropriate. Ancestry-specific fixed-effects inverse-variance weighted meta-analyses of study-specific results, with genomic control correction, were conducted in METAL (http://www.sph.umich.edu/csg/abecasis/metal/). Meta-analyses included approximately 11.1 million variants for European ancestry, 18.1 million for African ancestry, 4.2 million variants for Asian ancestry, and 13.8 million for Hispanic/Latino ethnicity. See Methods for full methods description.

European ancestry meta-analysis identified 17 novel loci (defined as more than 500kb in either direction from a known locus) ^16, 17^ which were significantly (defined as p<5.0x10^−8^) ^14, 18^ associated with pulmonary function: 2 loci for FEV_1_ only, 6 loci for FVC only, 7 loci for FEV_1_/FVC only, and 2 loci for both FEV_1_ and FVC (Table 2, Figure 1, Supplemental Figures 2-3). The African ancestry meta-analysis identified 8 novel loci significantly associated with pulmonary function: 2 loci for FEV_1_, 1 locus for FVC, and 5 loci for FEV_1_/FVC (Table 2, Supplemental Figure 1-3). Five of these loci were also significant at a stricter p<2.5x10^−8^ threshold as has been suggested for populations of African ancestry^18^. Six of the African ancestry loci were identified based on low frequency variants (allele frequencies 0.01 to 0.02). In the Hispanic/Latino ethnicity meta-analysis, we identified one novel locus for FVC (Table 2, Supplemental Figure 1-3). Another locus was significantly associated with FEV_1_, but this region was recently reported by HCHS/SOL^19^. For FEV_1_/FVC among Hispanics/Latinos, all significant variants were in loci identified in previous studies of European ancestry populations. In the Asian ancestry meta-analysis, all variants significantly associated with PFTs were also in loci previously identified among European ancestry populations (Supplemental Figure 1). Within each ancestry, variants discovered for one PFT were also looked-up for associations with the other two PFTs (Supplemental Table 1).

**Table 2.**
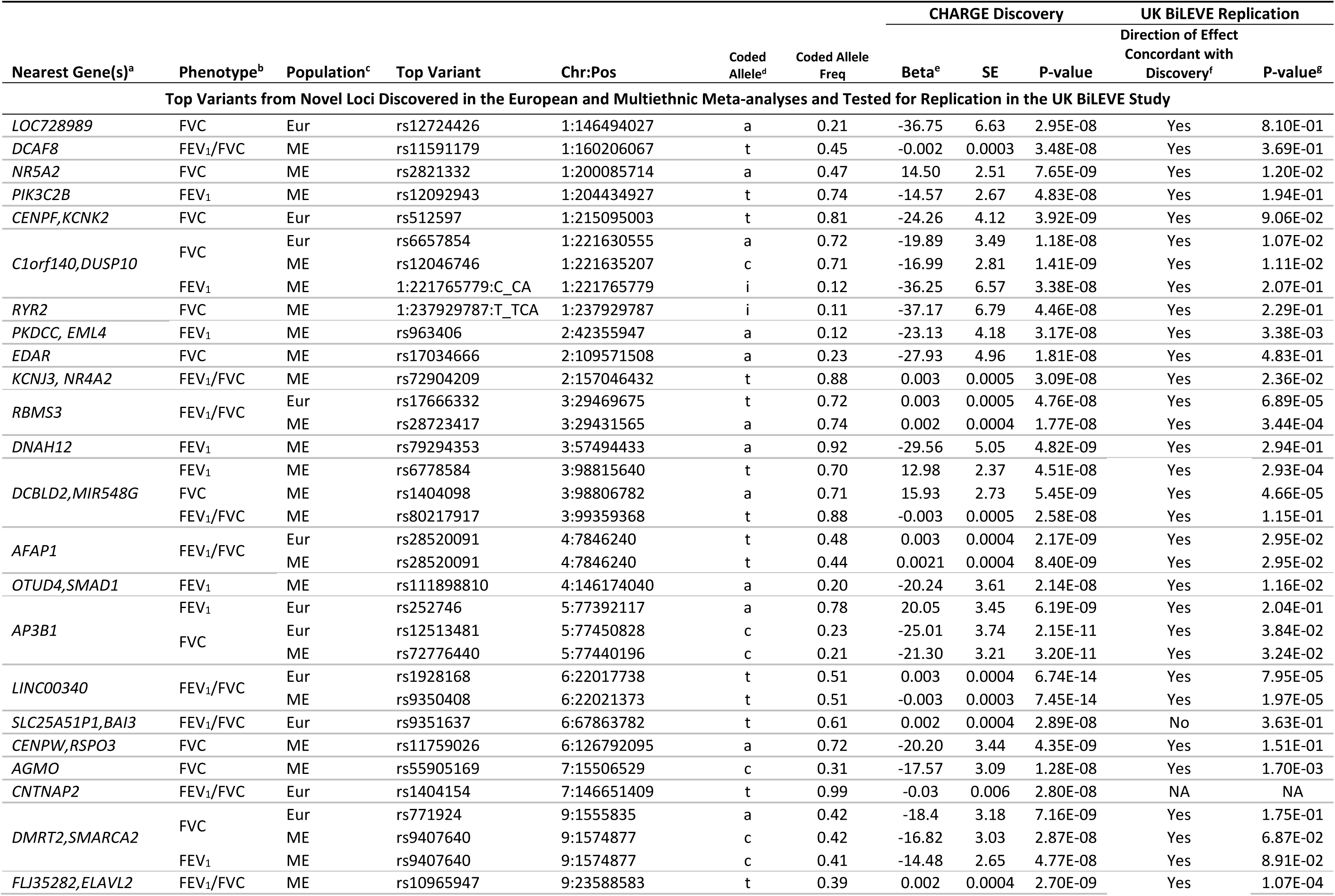
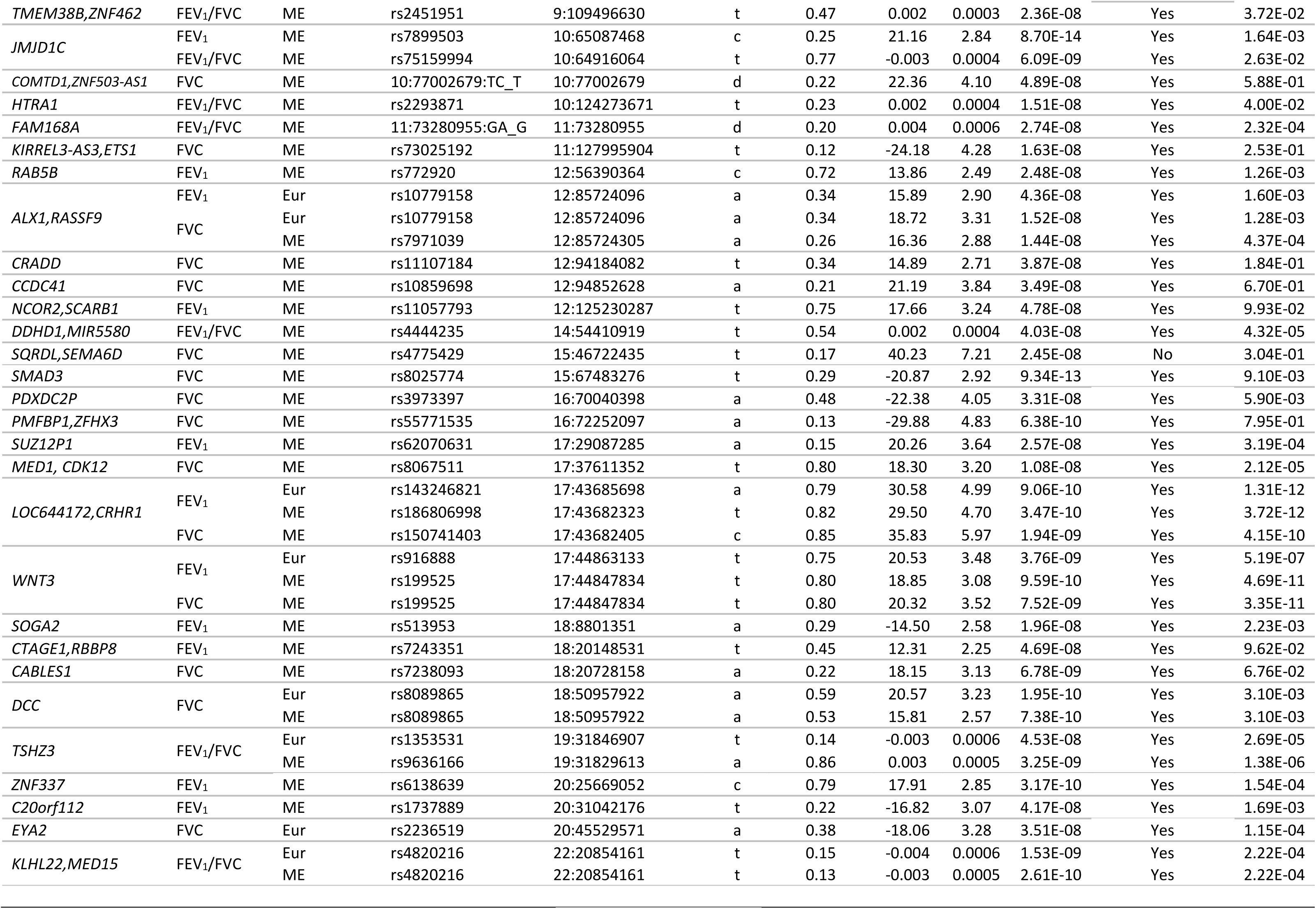
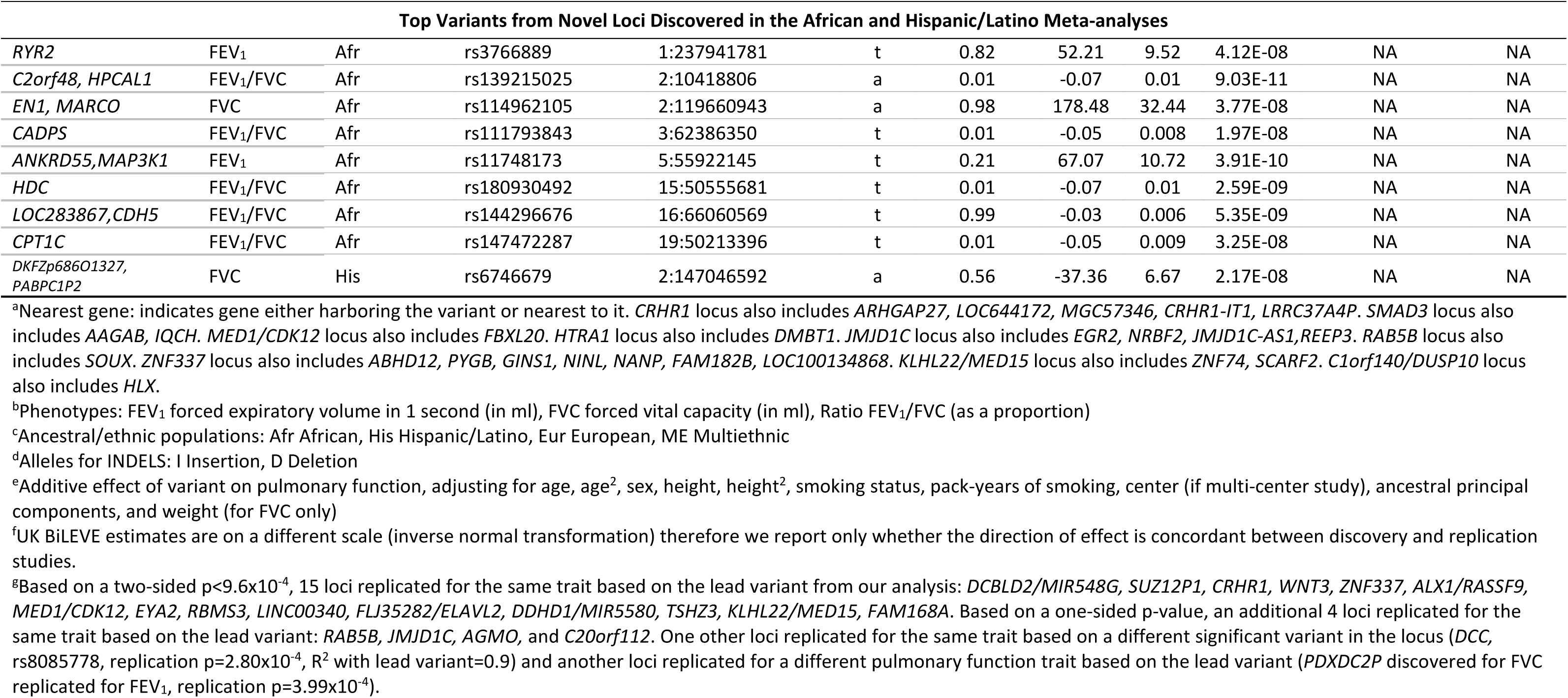
Top Variants from Novel Loci Discovered in the CHARGE Consortium Meta-Analysis of Pulmonary Function

**Figure 1.**
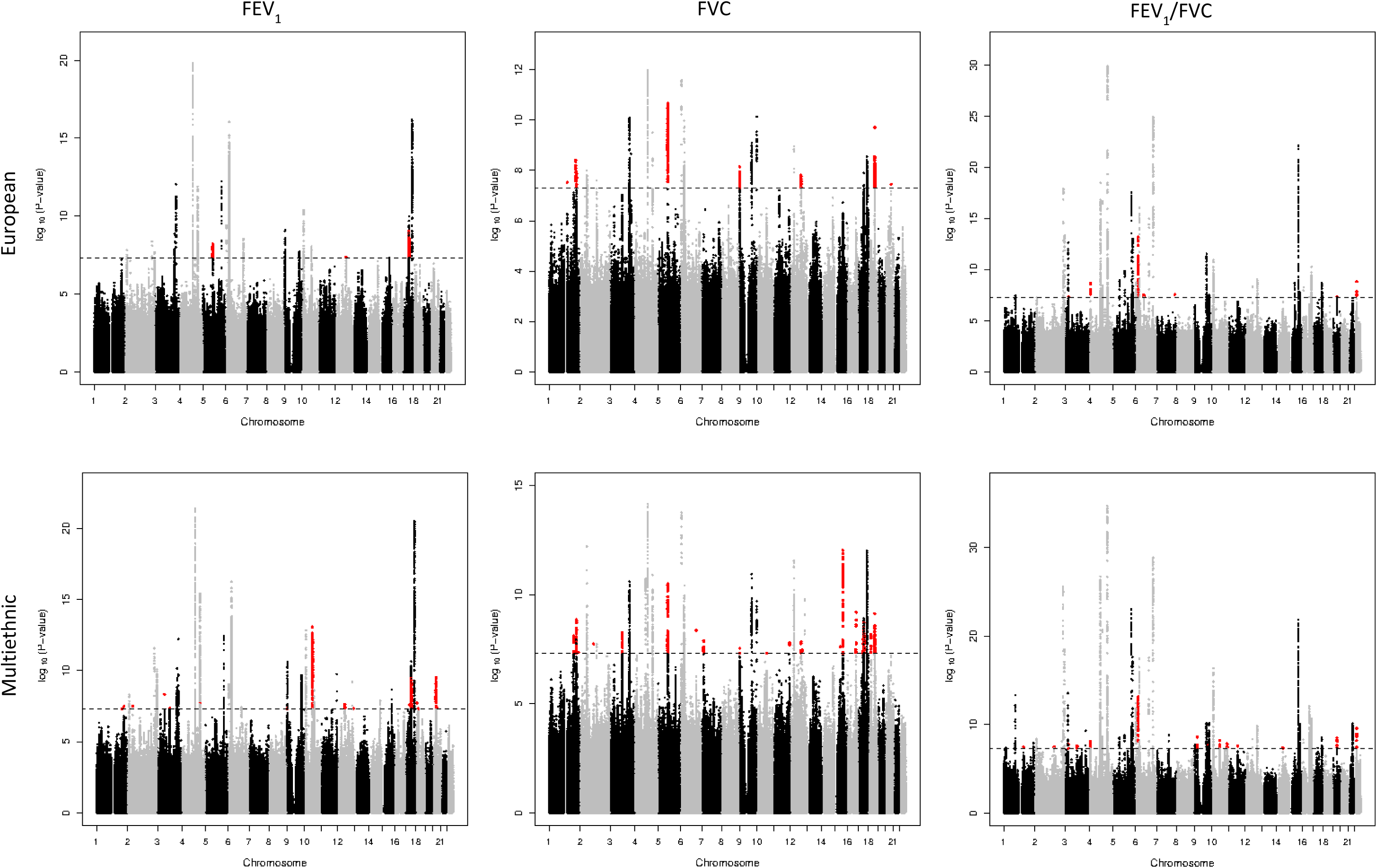
Manhattan plots for genome-wide association results for pulmonary function (FEV_1_, FVC, and FEV_1_/FVC) from European and multiethnic meta-analyses in CHARGE. Novel loci indicated by red. Significance level (5x10^−8^) indicated by dashed line.

### Multiethnic Meta-analysis

In multiethnic fixed effects meta-analyses of 10.9 million variants, we identified 47 novel loci significantly associated with pulmonary function. Thirteen of these loci were also identified in the ancestry-specific meta-analyses, while 34 were uniquely identified in the multiethnic meta-analysis: 11 loci for FEV_1_ only, 14 loci for FVC only, 7 loci for FEV_1_/FVC only, 1 locus for FEV_1_ and FEV_1_/FVC, and 1 locus for all three phenotypes (Table 2, Figure 1, Supplemental Figure 2-3). Although many of the 34 loci uniquely identified in the multiethnic meta-analysis were just shy of significance in the European ancestry meta-analysis, and therefore benefited from the additional sample size of the multiethnic meta-analysis, some multiethnic loci contained variants near genome-wide significance in at least one other ancestry-specific meta-analysis with some nominal significance (p<0.05) in the remaining ancestry-specific meta-analyses (Supplemental Table 2). For example, rs7899503 in *JMJD1C* associated with FEV_1_ had an *I*^2^ value of 0 with a heterogeneity p-value of 0.40 and meta-analysis p-values of 1.35x10^−5^ for European ancestry, 4.56x10^−7^ for Asian ancestry, 0.002 for Hispanic/Latino ethnicity and 0.03 for African ancestry, with a multiethnic p-value of 8.70x10^−14^.

In addition to the fixed-effects multiethnic meta-analysis, we conducted a random-effects meta-analysis using the Han and Eskin method^20^ in METASOFT (http://genetics.cs.ucla.edu/meta/) as a sensitivity analysis. In instances where significant heterogeneity is present, the Han-Eskin method mitigates power loss^20^. In the Han-Eskin random-effects model, 37 of the 47 loci identified in the fixed-effects model at p<5x10^−8^ had a p-value below the same threshold (Supplemental Table 3). Among the 10 loci that did not, 8 loci still gave a p<5x10^−7^ in the Han-Eskin random-effects model (*PIK3C2B, SUZ12P1, NCOR2/SCARB1, CTAGE1/RBBP8, C20orf112, COMTD1/ZNF503-AS1, EDAR,* and *RBMS3*) while only two did not (*CRADD* and *CCDC41*) (Supplemental Table 3). In addition, there were 6 loci for FEV_1_/FVC that were genome-wide significant in the Han-Eskin random-effects model that had not quite achieved genome-wide significance in the fixed-effects model: G*STO1/GSTO2* (chr10, rs10883990), *FRMD4A* (chr10, rs1418884), *ETFA/SCAPER* (chr15, rs12440815), *APP* (chr21, rs2830155), *A4GNT* (chr3, rs9864090), *UBASH3B* (chr11, rs4935813) (Supplemental Table 3).

### X-Chromosome Meta-Analysis

Within 13 of the 22 studies we conducted linear regression to model the effects of X-chromosome variants on PFTs by sex. Fixed-effects inverse-variance weighted meta-analyses were conducted separately in males and females using METAL and the resulting sex-specific results were combined using a weighted sums approach. No X-chromosome variants were associated with PFTs at genome-wide significance in any ancestral population (data not shown).

### Look-up Replication of Novel Loci

Look-up replication was conducted in the UK BiLEVE study (N=48,943)^14^. Since this study only included individuals of European ancestry, we sought replication only for the 52 novel loci (excluding the major histocompatibility complex, MHC) identified in either the European ancestry or multiethnic discovery meta-analyses. Data for the lead variant was available in the UK BiLEVE study for 51 loci, including 49 loci with a consistent direction of effect between our results and those from UK BiLEVE (Table 2). Based on a two-sided p<9.6x10^−4^ (0.05/52), 15 loci replicated for the same trait based on the lead variant from our analysis: *DCBLD2/MIR548G, SUZ12P1, CRHR1, WNT3, ZNF337, ALX1/RASSF9, MED1/CDK12, EYA2, RBMS3, LINC00340, FLJ35282/ELAVL2, DDHD1/MIR5580, TSHZ3, KLHL22/MED15, FAM168A* (Table 2). It was recently demonstrated that using one-sided replication p-values in GWAS, guided by the direction of association in the discovery study, increases replication power while being protective against type 1 error compared to the two-sided p-values^21^; under this criterion, an additional 4 loci replicated for the same trait based on the lead variant: *RAB5B, JMJD1C, AGMO,* and *C20orf112* (Table 2). Finally, one locus replicated for the same trait based on a different significant variant in the locus *(DCC,* rs8085778, replication p=2.80x10^−4^, R^2^ with lead variant=0.9) and one locus replicated for a different pulmonary function trait based on the lead variant (*PDXDC2P,* rs3973397, discovered for FVC replicated for FEV_1_, replication p=3.99x10^−4^). In summary, we found evidence of replication in UK BiLEVE for 21 novel loci.

### Overlap of Newly Identified Lung Function Loci with COPD

Pulmonary function measures are the basis for the diagnosis of COPD, an important clinical outcome; therefore, we also looked-up the 52 novel loci identified in the European ancestry or multiethnic meta-analyses in the International COPD Genetics Consortium (ICGC). This consortium recently published a meta-analysis of 1000 Genomes imputed variants and COPD primarily among individuals of European ancestry (N= 15,256 cases and 47,936 controls), including some of the same individuals included in the present lung function analysis^22^. Ten lead variants representing 8 novel loci were associated with COPD at p<9.6x10^−4^: *RBMS3, OTUD4/SMAD1, TMEM38B/ZNF462, NCOR2/SCARB1, SUZ12P1, WNT3, SOGA2, C20orf112* (Supplemental Table 4). Directions of effects were consistent between our results and the COPD findings for these variants (i.e. variants associated with increased pulmonary function values were associated with decreased odds of COPD and vice-versa). Our top variant in *SOGA2* (also known as *MTCL1*) is in LD (R^2^=0.8) with the top variant for COPD as reported by the IGCG Consortium^22^.

### Expression Quantitative Trait Loci (eQTL) and Methylation Quantitative Trait Loci (mQTL) Signals of Pulmonary Function Variants

To query whether novel loci contained variants associated with gene expression (eQTLs), we looked-up variants from all 60 novel loci identified in ancestry-specific or multiethnic meta-analyses in the following datasets: 1) lung samples from 278 individuals in GTEx^23^; 2) lung samples from 1,111 individuals participating in studies from the Lung eQTL Consortium including Laval University, the University of Groningen and the University of British Columbia^24-26^; 3) whole blood samples from 5,257 individuals participating in the Framingham Heart Study^27^; 4) peripheral blood samples from 5,311 individuals participating in studies from EGCUT, InCHIANTI, Rotterdam Study, Fehrmann, HVH, SHIP-TREND and DILGOM^28^; and 5) peripheral blood samples from 2,116 individuals participating in 4 Dutch studies collectively known as BIOS^29, 30^. The Lung eQTL Consortium study used a 10% FDR cut-off, while all other studies used a 5% FDR cut-off (see Supplemental Methods for further study descriptions and methods).

A significant cis-eQTL in at least one dataset was found for 34 lead variants representing 27 novel loci (Supplemental Table 5). Of these, 13 loci had significant cis-eQTLs only in datasets with blood samples and 3 loci only in datasets with lung samples (*COMTD1/ZNF503-AS1, FAM168A, SOGA2*). Eleven loci had significant cis-eQTLs in both blood and lung samples based on lead variants, with 1 locus having a significant cis-eQTL across all five datasets (*SMAD3*) and another 4 loci having a significant cis-eQTL in four datasets (*RAB5B, CRHR1, WNT3, ZNF337*). Significant trans-eQTLs in at least one dataset were found for 7 lead variants representing 4 novel loci (*TMEM38B/ZNF462*, *RAB5B, CRHR1*, and *WNT3*, Supplemental Table 5).

In addition, mQTL data were available from FHS and BIOS. Significant cis-mQTLs and trans-mQTLs in at least one dataset were found for 52 lead variants (43 novel loci) and 1 lead variant (1 novel locus), respectively (Supplemental Table 5).

### Heritability and Genetic Correlation with Smoking and Height (LD Score Regression)

We used LD score regression^31^ to estimate the variance explained by genetic variants investigated in our European ancestry meta-analysis, also known as SNP heritability. Across the genome, the SNP heritability (narrow-sense) was estimated to be 20.7% (SE 1.5%) for FEV_1_, 19.9% (SE 1.4%) for FVC and 17.5% (SE 1.4%) for FEV_1_/FVC.

We also partitioned heritability by functional categories to investigate whether particular subsets of common variants were enriched^32^. We found significant enrichment in 6 functional categories for all three PFTs: conserved regions in mammals, DNase I hypersensitive sites (DHS), super enhancers, the histone methylation mark H3K4me1 and histone acetylation marks H3K9Ac and H3K27Ac (Supplemental Figure 4). Another 7 categories showed enrichment for at least one PFT (Supplemental Figure 5). We observed the largest enrichment of heritability (14.5-15.3 fold) for conserved regions in mammals, which has ranked highest in previous partitioned heritability analyses for other GWAS traits (Supplemental Figure 5)^32^.

Since both height and smoking are important determinants of pulmonary function, and have been associated with many common variants in previous GWAS, we also used LD score regression to investigate genetic overlap^33^ between our FEV_1_, FVC and FEV_1_/FVC results and publicly available GWAS results of ever smoking^34^ and height^35^. No significant genetic correlation was found between PFTs and smoking or height (Supplemental Table 6), indicating our findings are independent of those traits.

### Functional Annotation of Pulmonary Function Variants

For functional annotation, we considered all novel variants with p<5x10^−8^ from the 60 loci discovered in our ancestry-specific and multiethnic meta-analyses, plus significant variants from the MHC region, two loci previously discovered in the CHARGE exome chip study (*LY86/RREB1* and *SEC24C*)^36^ and *DDX1*. Using Ensembl VEP^37^, we found 6 missense variants in 4 loci outside of the MHC region and 22 missense variants in the MHC region (Supplementary Table 7). Of the 28 total missense variants, two (chr15:67528374 in *AAGAB* and chr6:30899524 in the MHC region) appear to be possibly damaging based on SIFT^38^ and PolyPhen-2^39^ scores (Supplementary Table 7). Using CADD^40^, we found an additional 28 deleterious variants from 15 loci based on having a scaled C-score greater than 15 (Supplementary Table 8). In the MHC region, we found another 11 deleterious variants based on CADD. Based on RegulomeDB^41^, which annotates regulatory elements especially for non-coding regions, there were 52 variants from 18 loci with predicted regulatory functions (Supplementary Table 8). In the MHC region, there were an additional 72 variants with predicted regulatory functions.

### Pathway Enrichment Analysis (DEPICT and IPA)

Gene set enrichment analyses conducted using DEPICT^42^ on genes annotated to variants with p<1x10^−5^ based on the European ancestry meta-analysis results revealed 218 significantly enriched pathways (FDR<0.05) (Supplementary Table 9). The enriched pathways were dominated by fundamental developmental processes, including many involved in morphogenesis of the heart, vasculature, and lung. Tissue and cell type analysis noted significant enrichment (FDR<0.05) of smooth muscle, an important component of the lung (Supplemental Table 10 and Supplemental Figure 6). We found 8, 1, and 82 significantly prioritized genes (FDR<0.05) for FEV_1_, FVC, and FEV_1_/FVC, respectively (Supplemental Table 11). Given the generally smaller p-values for variants associated with FEV_1_/FVC, enriched pathways and tissue/cell types as well as prioritized genes were predominantly discovered from DEPICT analyses of FEV_1_/FVC.

Due to extended LD across the MHC locus on chromosome 6 (positions 25,000,000-35,000,000), DEPICT excludes this region^42^. Standard Ingenuity Pathway Analysis (IPA) run without excluding the MHC highlighted 16 enriched networks, including three involved in inflammatory diseases or immunity; 33 of the 84 genes in these three networks are in the MHC region (Supplementary Table 12).

### Identification of Potential Causal Variants by Incorporating Trans-ethnic Data and Functional Annotation (PAINTOR)

Using a multiethnic fine-mapping analysis incorporating strength of association, variation in genetic background across major ethnic groups, and functional annotations in PAINTOR^43^, we examined 40 loci that contained at least 5 genome-wide significant variants in the European ancestry and multiethnic meta-analyses or at least 1 significant variant in the African or Hispanic/Latino ancestry meta-analyses. We identified 15 variants representing 13 loci as having high posterior probabilities of causality (>0.8): 3 for FEV_1_, 3 for FVC, and 9 for FEV_1_/FVC (Supplemental Table 13). Of the 15 putative casual variants, 11 showed high posterior probabilities of causality (>0.8) before considering annotations and 4 were identified by adding functional annotations. Nine were the top SNPs at that locus from the fixed-effects meta-analysis (loci: *WNT3*, *PMFBP1/ZFHX3*, *EN1/MARCO*, *C2orf48/HPCAL1*, *CPT1C*, *CADPS*, *LOC283867/CDH5*, *HDC*, and *CDC7/TGFBR3*), while 6 were not (loci: *CDK2/RAB5B*, *BMS1P4*, *PMFBP1/ZFHX3*, *FLJ35282/ELAVL2*, *HDC*, and *COL8A1*).

### Identification of Independent Signals in Known and Novel Loci for Pulmonary Function (FINEMAP)

We used FINEMAP^44^ to identify variants with a high posterior probability of causality (>0.6) independent of 118 lead variants in pulmonary function loci identified in the current or previous studies^14^. We identified 37 independent variants for 23 previously identified loci and one independent variant at each of two novel loci (*LINC00340* and *SLC25A51P1/BAI3*; Supplementary Table 14).

### Gene-based Analysis of GWAS Summary Statistics (S-PrediXcan)

Among the novel loci identified in the current GWAS of PFTs, we identified 7 variants corresponding to 9 genes demonstrating genome-wide significant evidence of association with lung or whole blood tissue-specific expression (Supplemental Table 15) based on the gene-based S-PrediXcan approach^45^. Bayesian colocalization analysis^46^ indicated the following associations demonstrated at least 50% probability of shared SNPs underlying both gene expression and PFTs: *ARHGEF17* and *FAM168A* in analysis of multiethnic GWAS for FEV_1_/FVC based on GTEx whole blood models, and *WNT3* in analysis of multiethnic GWAS for FVC based on GTEx lung models (Supplemental Table 16).

### Druggable Targets

To investigate whether the genes identified in our study encode proteins with predicted drug targets, we queried the ChEMBL database (https://www.ebi.ac.uk/chembl/). In addition, we incorporated an Ingenuity Pathway Analysis (IPA) to identify potential upstream targets. Genes associated with pulmonary function, but not included in the drug target analysis performed by Wain et al^14^, were evaluated for a total of 139 genes outside of the MHC: 110 genes representing the 60 novel loci identified in our fixed-effects ancestry-specific and multiethnic meta-analysis, 13 genes representing the 6 novel loci identified in our random effects meta-analysis^20^, 3 genes representing an additional 3 loci near significance in the African ancestry meta-analysis (*BAZ2B, NONE/PCDH10, and ADAMTS17*), 9 genes representing 2 loci identified in a recent CHARGE analysis of exome variants^36^ which were also significant in our 1000 Genomes analysis (*LY86/RREB1* and *SEC24C*), and 4 genes representing one locus identified at genome-wide significance in a separate publication from one of our participating studies (HCHS/SOL)^19^ but also significant in our analysis (*ADORA2B/ZSWIM7/TTC19/NCOR1*). In the ChEMBL database, 17 of these genes encode proteins with predicted or known drug targets: *NR5A2, KCNK2, EDAR, KCNJ3, NR4A2, BAZ2B, A4GNT, GSTO1, GSTO2, NCOR2, SMAD3, NCOR1, CDK12, WNT3, PYGB, NANP, EYA2* (Supplemental Table 17). Of these, two genes (*KCNK2 and CDK12)* have approved drug targets. Using IPA, four additional genes were predicted as drug targets (*ADORA2B, APP, CRHR1, and MAP3K1*; Supplemental Table 18) and 37 genes had drugs or chemicals as upstream regulators (Supplemental Table 19).

## Discussion

By conducting a GWAS meta-analysis in a large multiethnic population we increased the number of known loci associated with pulmonary function by over 50%. In total, we identified 60 novel genetic regions (outside of the MHC region): 17 from European ancestry, 8 from African ancestry, 1 from Hispanic/Latino ethnicity, and 34 from multiethnic meta-analyses.

Just under half of the novel loci identified in our European ancestry and multiethnic meta-analyses replicated in look-up in a smaller independent sample of Europeans from the UK BiLEVE study^14^: *DCBLD2/MIR548G, SUZ12P1, CRHR1, WNT3, ZNF337, ALX1/RASSF9, MED1/CDK12, EYA2, RBMS3, LINC00340, FLJ35282/ELAVL2, DDHD1/MIR5580, TSHZ3, KLHL22/MED15, FAM168A*, *RAB5B, JMJD1C, AGMO, C20orf112*, *DCC,* and *PDXDC2P*. Among those loci which did not directly replicate for PFTs in the UK BiLEVE, the lead variants in an additional 4 European or multiethnic loci were significantly associated in the ICGC Consortium with COPD, which was defined using PFT measures^22^: *OTUD4/SMAD1, TMEM38B/ZNF462, NCOR2/SCARB1*, and *SOGA2.* Since we were not able to seek external replication for loci identified in the African ancestry meta-analyses, we used PAINTOR to query whether variants in the African-specific loci had a high probability of causality based on association statistics, LD, and functional annotations across ancestral groups^43^. Interestingly, PAINTOR identified the lead variants for 6 out of 8 novel loci identified in African ancestry as having a high probability for causality: *C2orf48/HPCAL1, EN1/MARCO, CADPS, HDC, LOC283867/CDH5*, and *CPT1C*. PAINTOR or other new genomic methods such as FINEMAP and S-PrediXcan also produced evidence for causality for 4 European ancestry and multiethnic loci which had not replicated in UK BiLEVE or ICGC: *DCAF8, AFAP1*, *SLC25A51P1/BAI3* and *SMAD3*. Therefore, we found evidence for look-up replication of 25 loci in the UK BiLEVE study or ICGC COPD consortium and support for validation of an additional 10 loci using PAINTOR, FINEMAP, or S-PrediXcan.

Our analysis also sheds light on additional potential causal genes at a complex locus (chromosome 17 near positions 43600000 to 44300000, hg19) previously discovered from GWAS of FEV_1_ which identified *KANSL1* in European populations as the top finding for this region^14, 15^. With the exception of a single INDEL in *KANSL1* in our European ancestry meta-analysis (17:44173680:T_TC, p=1.03x10^−10^), we found *CRHR1* as the strongest gene associated with FEV_1_ in this region. Although some variants in *CRHR1* identified in our study are within 500kb of *KANSL1* (e.g., rs16940672, 17:43908152, p=2.07x10^−10^), a number of significant variants in this gene are more than 500kb away from previously identified hits [our definition of novel] (e.g., rs143246821, 17:43685698, p=9.06x10^−10^). In our multiethnic meta-analysis, several variants in *CRHR1* were associated with FEV_1_ at smaller p-values than variants in *KANSL1*. Definitive assessment of the causal variants at this locus, as well as other multigenic GWAS loci, will likely require additional data from ongoing large scale sequencing studies to enable detailed fine mapping.

In both our European and multiethnic meta-analyses we also noted a significant association with *WNT3* on chromosome 17 near position 44800000 (hg19) which is more than 500kb from *KANSL1* or *CRHR1* [our definition of novel]. We found that the top variant in *WNT3* for FEV_1_ among individuals of European ancestry (rs916888, 17:44863133, p=3.76x10^−9^) had a high probability for causality based on PAINTOR, an analysis which integrates functional annotations along with association statistics and LD for each ethnicity^43^. We also found evidence that *WNT3* may be the causal gene at this locus using S-PrediXcan, a gene level association test that prioritizes potentially causal genes while filtering out LD-induced false positives^45^,^46^. Notably, S-PrediXcan implicated *WNT3* as a likely mediating gene for FVC based on the top variant in our multiethnic meta-analyses (rs199525, 17:44847834, p=7.52x10^−9^), which is an eQTL SNP for *WNT3* in lung and other tissues. Further, the lead *WNT3* variants for both FEV_1_ and FVC (rs916888 and rs199525) were significantly associated with COPD in a look-up of a large published meta-analysis dataset^22^. In addition, other genes in the *WNT* signaling pathway, a fundamental development pathway, have been implicated as influencing pulmonary function^47^. This pathway was also one of the significant pathways identified in our analysis. In a previous pathway analysis of asthma, *SMAD3* has been shown to interact with the *WNT* signaling pathway^48^. Finally, *WNT3* also emerged as having a potential druggable target, and incorporation of pathway analysis to identify upstream regulators found an additional four drugs in clinical use for which *WNT3* is a target molecule (chemotherapeutic agents doxorubicin and paclitaxel, the hormone beta-estradiol and LGK-974, a novel agent that targets a WNT-specific acyltransferase)^49^. Again, further evaluation of this interesting and complex locus which contains many significant variants in LD will benefit from data being generated in ongoing large-scale sequencing studies.

Some genes identified in our study play key roles in inflammation, immunity and pulmonary biology. For example, *MARCO* (macrophage receptor with collagenous structure) has been shown in murine models to be required for lung defense against pneumonia as well as inhaled particles^50^,^51^. *SMAD3* is part of the SMAD family of proteins which are signal transducers and transcriptional modulators that mediate multiple signaling pathways. *SMAD3* is activated by transforming growth factor beta (TGF-B) which plays a key role in airway remodeling. S*MAD3* has a predicted drug target and SNPs in *SMAD3* are significantly associated with asthma in GWAS^52^,^53^.

Other genes identified in our study which are targeted by approved drugs include *CDK12* and *KCNK2*. *CDK12* drug targets include AT-7519, Roniciclib, AZD-5438, and PH.A-793887. Roniciclib has been used in clinical trials including lung cancer patients^54^. *KCNK2* (potassium channel subfamily K member 2) is targeted by 5 inhalational anesthetic agents. These agents have anti-inflammatory effects both systemically^55^ and in the lungs^56^ and meta-analysis of clinical studies shows protection against pulmonary complications after cardiac surgery^57^. A recent trial suggested that one of these inhalation agents, sevoflurane, offers promise for reducing epithelial injury and improving outcomes in patients with acute respiratory distress syndrome^58^.

In addition to querying commonly used genome databases for functional annotation of variants, we sought to narrow down causal variants in implicated loci using recently developed methods that incorporate LD, functional data and/or the multiethnic analysis done in this paper. In particular, PAINTOR is a useful tool to identify potential causal variants in our novel loci as it leverages LD across ancestral groups along with association statistics and functional annotations^43^. PAINTOR identified 15 putative causal variants from 13 loci, including 6 novel loci in African ancestry meta-analyses and 7 loci uniquely identified in the multiethnic meta-analyses such as *PMFBP1/ZFHX3* and *COL8A1* (part of the *DCBLD2* loci). Notably, 8 of the 15 putative causal variants from PAINTOR were the top SNPs from the fixed-effects meta-analysis (e.g., rs916888 *WNT3*). Similarly, FINEMAP has been shown to be an accurate and efficient tool for investigating whether lead SNPs for a given loci are driven by independent variants in the same region, especially when annotation information is not available^44^. Among previous and novel loci identified in individuals of European ancestry, we identified 37 independent variants for 23 previously identified loci and two lead variants for two novel loci (rs1928168 *LINC00340* and rs9351637 *SLC25A51P1/BAI3)* with a high probability of causality. Finally, we ran S-PrediXcan a gene level association test that prioritizes potentially causal genes^45^. Seven of our novel loci contained putative causal genes based on S-PrediXcan for lung or whole blood tissues, including *NRBF2* (part of the *JMJD1C* locus) and *WNT3*. S-PrediXcan also highlighted the region around chromosome 11 position 73280000 (hg19), noting strong evidence for both *FAM168A* and *ARHGEF17* which was further supported by the co-localization analysis. Interestingly, DEPICT also prioritized *ARHGEF17*, a member of the guanine nucleotide exchange factor (GEF) family of genes which can mediate actin polymerization and contractile sensitization in airway smooth muscle^59^,^60^.

Rather than conducting a standard gene-based pathway analysis, we performed a newer integrative method, DEPICT, that incorporates cell and tissue-specific functional data into a pathway analysis to prioritize genes within implicated loci^42^. In addition to identifying potential causal variants, this approach revealed a number of fundamental development processes, including pathways related to lung development, regulation of growth, and organ morphogenesis. The *WNT* signaling pathway was also highlighted along with processes relevant to the pathogenesis of COPD including extracellular matrix structure and collagen networks. Tissue/cell type enrichment results highlighted smooth muscle which is highly relevant for lung function. DEPICT excludes the MHC due to extended LD in this region, which likely explains the relative paucity of inflammation-related pathways identified compared to previous pathway analyses in GWAS of PFTs^24^,^47^. Indeed, standard IPA analysis of our data including the MHC region, found that 33 of 84 genes (39%) in the 3 (out of 16) enriched networks involved in immune or inflammatory processes are in the MHC. The predominance of fundamental pathways related to lung growth, differentiation and structure is consistent with recent work^61^ that has rekindled interest in the observation made 40 years ago^62^ that individuals can cross the threshold for diagnosis of COPD either by rapid decline in adulthood or by starting from a lower baseline of maximal pulmonary function attained during growth. Within this context, understanding the genetic (and environmental) factors that influence the variability in maximal lung function attained during the first three decades of life is essential to reducing the public health burden of COPD^63^.

In summary, our study extends existing knowledge of the genetic landscape of PFTs by utilizing the more comprehensive 1000 Genomes imputed variants, increasing the sample size, including multiple ancestries and ethnicities, and employing newly developed computational applications to interrogate implicated loci. We discovered 60 novel loci associated with pulmonary function and replicated many in an independent sample. We found evidence that several variants in these loci were missense mutations and had possible deleterious or regulatory effects, and many had significant eQTLs. Further, using new genomic methods that incorporate LD, functional data and the multiethnic structure of our data, we shed light on potential causal genes and variants in implicated loci. Finally, several of the newly identified genes linked to lung function are druggable targets, highlighting the clinical relevance of our integrative genomics approach.

## Methods

### Studies

Member and affiliate studies from The <u>C</u>ohorts for <u>H</u>eart and <u>A</u>ging <u>R</u>esearch in <u>G</u>enomic <u>E</u>pidemiology (CHARGE) consortium with pulmonary function and 1,000 Genomes imputed genetic data were invited to participate in the present meta-analysis. Participating studies included: AGES, ALHS, ARIC, CARDIA, CHS, FamHS, FHS, GOYA, HCHS/SOL, HCS, Health ABC, Healthy Twin, JHS, KARE3, LifeLines, LLFS, MESA, NEO, 1982 PELOTAS, RSI, RSII, RIII. Characteristics of these studies are provided in Supplemental Table 20 and descriptions of study designs are provided in the Supplemental Methods.

### Pulmonary Function

Spirometry measures of pulmonary function (FEV_1_, FVC, and the ratio FEV_1_/FVC) were collected by trained staff in each study according to American Thoracic Society (ATS) or European Respiratory Society guidelines (see cohort descriptions in Supplemental Methods for more details).

### Variants

Studies used various genotyping platforms, including Affymetrix Human Array 6.0, Illumina Human Omni Chip 2.5, and others, as described in cohort descriptions in the Supplemental Methods. Using MACH, MINIMAC, or IMPUTE2, studies then used genotyped data to impute ∼38 million variants based on the 1000 Genomes Integrated phase 1 version 3 reference panel (released March 2012). Two studies (Hunter Community and CARDIA) imputed to the 1000 Genomes European phase 1 version 3 reference panel; sensitivity analyses excluding these two studies showed no material differences in results (see Supplemental Results).

### Statistical Analysis

Within each study, linear regression was used to model the additive effect of variants on PFTs. FEV_1_ and FVC were modeled as milliliters and FEV_1_/FVC as a proportion. Studies were asked to adjust analyses for age, age^2^, sex, height, height^2^, smoking status (never, former, current), pack-years of smoking, center (if multicenter study), and ancestral principal components, including a random familial effect to account for family relatedness when appropriate^64^. Models of FVC were additionally adjusted for weight. Analyses were conducted using ProbAbel, PLINK, FAST, or the R kinship package as described in the cohort descriptions of the Supplemental Methods.

Ancestry-specific and multiethnic fixed effects meta-analyses using inverse variance weighting of study-specific results with genomic control correction were conducted in Meta Analysis Helper (METAL, http://www.sph.umich.edu/csg/abecasis/metal/). Multiethnic random effects meta-analyses using the four ancestry-specific fixed effects meta-analysis results were conducted using the Han-Eskin model^20^ in METASOFT (http://genetics.cs.ucla.edu/meta/). Only variants with p-values for association <0.05 or p-values for heterogeneity <0.1 from the fixed-effects model were included in the random-effects model.

Variants with imputation quality scores (r^2^) less than 0.3 and/or a minor allele count (MAC) less than 20 were excluded from each study prior to meta-analysis. Following meta-analysis, we also excluded variants with less than 1/3 the total sample size or less than the sample size of the largest study for a given meta-analysis to achieve the following minimal sample sizes: 20,184 for European ancestry; 2,810 for African ancestry; 7,862 for Asian ancestry; 4,435 for Hispanic/Latino ethnicity and 30,238 for Multiethnic.

Significance was defined as p<5x10^−8^ ^14^,^18^. Novel variants were defined as being more than +/- 500kb from the top variant of a loci identified in a previous GWAS of pulmonary function^16^,^17^. We used the list of 97 known variants as published in the recent UK BiLEVE paper^14^ with the following modifications: added variants in *DDX1, DNER, CHRNA5* since listed in GWAS catalog; added variants in *LCT, FGF10, LY86/RREB1, SEC24C, RPAP1, CASC17*, and *UQCC1* since identified in exome chip paper^36^; added variant in *TMEM163* identified in Loth et al paper^10^; used 17:44339473 instead of 17:44192590 to represent *KANSL1* since 17:44339473 was the original variant listed for *KANSL1* in Wain et al 2015^15^; and used 12:28283187 instead of 12:28689514 to represent *PTHLH* since 12:28283187 was the original variants listed for *PTHLH* in Soler Artigas et al 2015^13^.

Genomic inflation factors (lambda values) from quantile-quantile plots of observed and expected p-values for ancestry- and phenotype-specific meta-analyses are presented in Supplemental Table 21. Lambda values were slightly higher in European and multiethnic meta-analyses (range of lambda 1.12 to 1.16) than in other ancestry-specific meta-analyses (range of lambda 1.01 to 1.06) likely due to the much larger sample sizes of the European and multiethnic meta-analyses^65^.

### LD Score Regression

The SNP heritability, i.e. the variance explained by genetic variants, was calculated from the European ancestry GWAS summary statistics (with genomic control off) using LD score regression (https://github.com/bulik/ldsc) ^31^. Partitioned heritability was also calculated using the method described by Finucane and colleagues^32^. In total, 28 functional annotation classes were used for this analysis, including: coding regions, regions conserved in mammals, CCCTC-binding factor (CTCF), DNase genomic foot printing (DGF), DNase I hypersensitive sites (DHS), fetal DHS, enhancer regions; including super-enhancers and active enhancers from the FANTOM5 panel of samples, histone marks including two versions of acetylation of histone H3 at lysine 27 (H3K27ac and H3K27ac2), histone marks monomethylation (H3K4me1), trimethylation of histone H3 at lysine 4 (H3K4me) and acetylation of histone H3 at lysine 9 (H3K9ac5). In addition to promotor and intronic regions, transcription factor binding site (TFBS), transcription start site (TSS) and untranslated regions (UTR3 and UTR5). A p-value of 0.05/28 classes < 1.79 x 10^−3^ was considered statistically significant. Genetic correlation between our pulmonary function (FEV_1_, FVC and FEV_1_/FVC) results and publicly available GWAS of ever smoking^34^ and height^35^ was also investigated using LD score regression^33^.

### Functional Annotation

To find functional elements in novel genome-wide significant signals, we annotated SNPs using various databases. We used Ensembl Variant Effect Predictor (VEP)^37^ (Accessed 17 Jan 2017) and obtained mapped genes, transcripts, consequence of variants on protein sequence, Sorting Intolerant from Tolerant (SIFT)^38^ scores, and Polymorphism Phenotyping v2 (PolyPhen-2)^39^ scores. We checked if there were deleterious variants using Combined Annotation Dependent Depletion (CADD) v1.3^40^ which integrates multiple annotations, compares each variant with possible substitutions across the human genome, ranks variants, and generates raw and scaled C-scores. A variant having a scaled C-score of 10 or 20 indicates that it is predicted to be in the top 10% or 1% deleterious changes in human genome, respectively. We used a cutoff of 15 to provide deleterious variants since it is the median for all possible splice site changes and non-synonymous variants (http://cadd.gs.washington.edu/info, Accessed 18 Jan 2017). To find potential regulatory variants, we used RegulomeDB^41^ (Accessed 17 Jan 2017) which integrates DNA features and regulatory information including DNAase hypersensitivity, transcription factor binding sites, promoter regions, chromatin states, eQTLs, and methylation signals based on multiple high-throughput datasets and assign a category to each variant. Variants having RegulomeDB categories 1 or 2, meaning ‘likely to affect binding and linked to expression of a gene target’ or ‘likely to affect binding,’ were considered as regulatory variants.

### Pathway Analysis (DEPICT and IPA)

For gene prioritization and identification of enriched pathways and tissues/cell types, we used Data-driven Expression Prioritized Integration for Complex Traits (DEPICT)^42^ with association results for FEV_1_, FVC, and FEV_1_/FVC. We used association results from our European ancestry meta-analysis and the LD structure from 1000 Genomes European (CEU, GBR, and TSI) reference panel. The software excludes the major histocompatibility complex (MHC) region on chromosome 6 due to extended LD structure in the region. We ran a version of DEPICT for 1000 Genomes imputed meta-analysis results using its default parameters with an input file containing chromosomal location and p-values for variants having unadjusted p-values <1x10^−5^. For gene set enrichment analyses, DEPICT utilizes 14,461 reconstituted gene sets generated by genes’ co-regulation patterns in 77,840 gene expression microarray data. For tissue/cell type enrichment analysis, mapped genes were tested if they are highly expressed in 209 medical subject headings (MeSH) annotations using 37,427 microarray data. Gene prioritization analysis using co-functionality of genes can provide candidate causal genes in associated loci even if the loci are poorly studied or a gene is not the closest gene to a genome-wide significant variant. We chose FDR<0.05 as a cutoff for statistical significance in these enrichment analyses and gene prioritization results. Because DEPICT excludes the MHC, we also ran a pathway analysis with Ingenuity Pathway Analysis (IPA) (Ingenuity Systems, Redwood City, CA, USA, http://www.ingenuity.com/) on genes to which variants with p<1x10^−5^ annotated.

### PAINTOR

To identify causal variants in novel genome-wide significant loci, we used a trans-ethnic functional fine mapping method^43^ implemented in PAINTOR (https://github.com/gkichaev/PAINTOR_FineMapping, Accessed on May 2, 2016). This method utilizes functional annotations along with association statistics (Z-scores) and linkage disequilibrium (LD) information for each locus for each ethnicity. We included our ancestry-specific meta-analysis results and used the African, American, European, and East Asian individuals from 1000 Genomes to calculate linkage disequilibrium (LD)^66^. From the novel loci we identified in our ancestry-specific and multiethnic fixed effects meta-analyses, we selected 40 high priority loci which had at least five variants meeting significance: 10 loci for FEV_1_, 15 loci for FVC, and 17 loci for FEV_1_/FVC. This list included 6 loci which overlapped with the UK BiLEVE 1000 Genomes paper^14^ and 1 locus with the CHARGE exome paper^36^, since we ran PAINTOR prior those publications. To reduce computational burden, we limited flanking regions to ±100 kilobase (kb) from the top single nucleotide polymorphisms (SNPs) and included variants with absolute value of Z-score greater than 1.96.

We used 269 publicly available annotations relevant to ‘lung’, ‘bronch’, or ‘pulmo’ from the following: hypersensitivity sites (DHSs)^67^, super enhancers^68^, Fantom5 enhancer and transcription start site regions^69^, immune cell enhancers^70^, and methylation and acetylation marks ENCODE^71^. We ran PAINTOR for each phenotype separately to prioritize annotations based on likelihood-ratio statistics^72, 73^. We included minimally correlated top annotations (less than five for each phenotype) to identify causal variants.

For the 40 loci from the fixed-effects meta-analysis, we used PAINTOR to construct credible sets of causal variants using a Bayesian meta-analysis framework. To obtain 95% credible sets for each locus, we ranked SNPs based on posterior probabilities of causality (high to low) and then took the SNPs filling in 95% of the summed posterior probability. We computed the median number of SNPs in the credible sets for ancestry-specific and multi-ethnic analyses of each trait.

### FINEMAP

We used FINEMAP^44^ to identify signals independent of lead variants for pulmonary function loci identified in the current or previous studies^14^. We used a reference population from the Rotterdam Study (N=6,291). SNPs with MAF of <1% were excluded, leaving 118 SNPs for analysis. 10 SNPs for FEV_1_ and FVC and 20 SNPs for FEV_1_/FVC were further excluded because the LD matrix of the reference file from the Rotterdam Study did not represent the correlation matrix of the total study population. We allowed up to 10 causal SNPs per loci in FINEMAP analyses. To reduce the chance of false positive findings, we also conducted sensitivity analyses allowing up to 15 causal SNPs for loci with more than 4 SNPs with posterior probabilities of >0.8.

### S-PrediXcan

S-PrediXcan is a novel summary statistics based approach for gene-based analysis^45^ that was derived as an extension of the PrediXcan method for integration of GWAS and reference transcriptome data^74^. We used the S-PrediXcan approach to prioritize potentially causal genes, coupled with a Bayesian colocalization procedure^46^ used to filter out LD-induced false positives. S-PrediXcan was used to analyze both European ancestry and multi-ethnic GWAS summary data for pulmonary function traits from the current study.

S-PrediXcan analysis was performed using the following publicly available tissue-specific expression models (http://predictdb.org) from the Genotype-Tissue Expression (GTEx) project v6p^23^: (1)GTEx Lung (278 samples) and (2) GTEx Whole blood (338 samples). Approximately 85% of participants in GTEx are white, 12% African American, and 3% of other races/ethnicities. Gene-based S-PrediXcan results were filtered on the following: (1) Proportion of SNPs used = (n SNPs available in GWAS summary data)/ (n SNPs in prediction model) > 0.6, and (2) prediction performance R-squared > 0.01. Following application of S-PrediXcan to each of the GWAS summary data sets, we computed Bonferroni-corrected p-values derived as the nominal p-value for each gene-based test divided by the number of genes passing specified filters in each analysis to test whether genetically regulated gene expression was associated with the trait of interest. The genome-wide S-PrediXcan results were then merged with novel loci from the current GWAS study by identifying all matches in which the novel locus SNP was within 500kb of the start of the gene.

We further incorporated a Bayesian colocalization approach^46^ to interpret the extent to which S-PrediXcan results may have been influenced by linkage disequilibrium within the region of interest. The Bayesian colocalization procedure was run using the following priors: p1 = 1e-4; prior probability SNP associated to trait 1, p2 = 1e-4; prior probability SNP associated to trait 2, p12 = 1e-5; prior probability SNP associated to both traits. The procedure generated posterior probabilities that correspond to one of the following hypotheses: a region is (H0) has no association with neither trait, (H1) associated with PFT phenotype but not gene expression, (H2) associated with gene expression but not PFT phenotype, (H3) associated with both traits, due to two independent SNPs, (H4) associated with both traits, due to one shared SNP.

### Druggable Targets

We searched annotated gene lists against the ChEMBL database (v22.1, updated on November 15, 2016) to identify genes as targets of approved drugs or drugs in development. In addition, we used the Ingenuity Pathway Analysis (IPA, www.ingenuity.com, content of 2017-06-22) to identify drug targets and upstream regulators of the gene lists. We reported the upstream regulators in the following categories, biologic drug, chemical - endogenous mammalian, chemical - kinase inhibitor, chemical – other, chemical drug, chemical reagent, and chemical toxicant.

### Data Availability

Upon acceptance at a journal, complete meta-analysis results will be deposited in the database of Genotypes and Phenotypes (dbGaP) under the CHARGE acquisition number phs000930. GWAS data for US studies are already available in dbGAP. For non-US studies, please send requests to the study PI or Stephanie London who will forward them to the relevant party.

## Author Contributions

Study-level Design and Data Collection: ABW, TS, MKL, LL, JCL, AVS, TMB, MFF, WG, TSA, WT, CO, QD, KdJ, MKW, XQW, RN, FPH, MG, TBH, RK, SRH, LP, KMB, YL, EGH, JGW, JMV, JS, RGB, RdM, AMBM, LJL, ACM, CMS, JCC, SBK, RJS, KC, JIR, TNB, FCW, NF, JAB, RCK, KL, MM, MAP, FRR, KDT, VG, KEN, MF, BMP, RHM, GO, TH, CCL, PC, JS, WJK, JRA, LL, HMB, BT, SSR, DOMK, BLH, AGU, GGB, SAG, JD, AM, SJL

Study-level Data Analysis: ABW, TS, MKL, LL, JCL, AVS, TMB, MFF, WG, TSA, WT, CO, QD, KdJ, MKW, XQW, RN, FPH

Meta-analysis: ABW, TS, JJ, JD, SJL

Pulmonary Function Look-up Replication in the UK BiLEVE Study: VEJ

COPD Look-up in the ICGC Consortium: BDH, MHC, International COPD Genetics Consortium Investigators

eQTL Look-up: ABW, LL, MO, TH, MvdB, RJ, YB, DCN, DDS

Functional Data Analysis (LD Score, DEPICT, PAINTOR, FINEMAP, S-PrediXcan, or Druggable Targets): MKL, NT, JNN, TW, GK, HHHA, HKI, AM

Writing Group: ABW, TS, MKL, NT, JNN, LL, GGB, SAB, JD, AM, SJL

Manuscript Review: all authors

### Acknowledgements

We thank Huiling Li for expert technical assistance and Dr. Frank Day for computational support, both from the National Institute of Environmental Health Sciences (NIEHS), and Dr. Louise Wain, University of Leicester, for critical reviews of the manuscript. Supported in part by the Intramural Research Program of the National Institutes of Health, NIEHS. Infrastructure for the CHARGE Consortium is supported in part by the National Heart, Lung, and Blood Institute grant R01HL105756. Study specific funding and acknowledgments can be found in the Supplemental Material.

